# Potassium rhythms couple the circadian clock to the cell cycle

**DOI:** 10.1101/2024.04.02.587153

**Authors:** Sergio Gil Rodríguez, Priya Crosby, Louise L. Hansen, Ellen Grünewald, Andrew D. Beale, Rebecca K. Spangler, Beverley M. Rabbitts, Carrie L. Partch, Alessandra Stangherlin, John S. O’Neill, Gerben van Ooijen

**Affiliations:** School of Biological Sciences, University of Edinburgh, Max Born Crescent EH9 3BF Edinburgh, United Kingdom; Department of Chemistry and Biochemistry, University of California Santa Cruz, Santa Cruz, CA, 95064, USA; UKRI MRC Laboratory of Molecular Biology, Francis Crick Ave, Cambridge, CB2 0QH, United Kingdom; Faculty of Medicine and University Hospital Cologne, Cluster of Excellence Cellular Stress Responses in Aging-associated Diseases (CECAD), Institute for Mitochondrial Diseases and Ageing, University of Cologne, Joseph-Stelzmann-Str, 50931, Cologne, Germany

## Abstract

Circadian (∼24 h) rhythms are a fundamental feature of life, and their disruption increases the risk of infectious diseases, metabolic disorders, and cancer^1–6^. Circadian rhythms couple to the cell cycle across eukaryotes^7,8^ but the underlying mechanism is unknown. We previously identified an evolutionarily conserved circadian oscillation in intracellular potassium concentration, [K^+^]_i_^9,10^. As critical events in the cell cycle are regulated by intracellular potassium^11,12^, an enticing hypothesis is that circadian rhythms in [K^+^]_i_ form the basis of this coupling. We used a minimal model cell, the alga *Ostreococcus tauri,* to uncover the role of potassium in linking these two cycles. We found direct reciprocal feedback between [K^+^]_i_ and circadian gene expression. Inhibition of proliferation by manipulating potassium rhythms was dependent on the phase of the circadian cycle. Furthermore, we observed a total inhibition of cell proliferation when circadian gene expression is inhibited. Strikingly, under these conditions a sudden enforced gradient of extracellular potassium was sufficient to induce a round of cell division. Finally, we provide evidence that interactions between potassium and circadian rhythms also influence proliferation in mammalian cells. These results establish circadian regulation of intracellular potassium levels as a primary factor coupling the cell- and circadian cycles across diverse organisms.

## Main

Evolution has provided most eukaryotes with an internal biological timekeeping system to anticipate predictable environmental changes that occur due to Earth’s daily rotation^13^. This endogenous and self-sustaining mechanism, colloquially known as the circadian clock, enhances physiology, metabolism, and overall fitness^13,14^. Conversely, altered or disrupted circadian rhythmicity impacts on many key cellular processes and results in increased risk of disorders and pathologies^15^, including metabolic syndrome and cancer^16–18^. At the cellular level, circadian rhythmicity involves the rhythmic expression of clock genes that engage in Transcriptional/Translational Feedback Loops (TTFLs)^13^, which is further regulated by highly conserved post-transcriptional mechanisms that are not currently fully understood^19,20^.

Bidirectional interaction between the circadian clock and the cell cycle has been described across organisms that display circadian rhythmicity, from cyanobacteria to mammals^21^. Cues that synchronise circadian rhythms with the external day/night cycle also shift timing of cell division and growth, indicating a preferred circadian time for cytokinesis^22–25^. Thus, while evidence exists of coupling between the circadian system and the cell cycle, the mechanistic connection has not been elucidated.

Previously, we reported circadian rhythms in the concentration of intracellular ions in representative species from all eukaryotic kingdoms^9^. These ion rhythms functionally regulate fundamental cellular processes including translation, metabolism, and proteostasis^9,10,26^. Potassium is the most abundant ion in any eukaryotic cell, constituting an average of 0.2% of the total body weight in humans^27^ and 2-10% of dry weight in plants^28^, and acts to maintain fluid and electrolyte balance over membranes^29^. Even small alterations in the intracellular balance and flux of potassium are linked to important metabolic and cell cycle disorders, including cancer^10,30^. Therefore, high-amplitude circadian rhythms in potassium likely impact upon crucial cellular processes such as membrane potential, ion homeostasis, enzyme activity, and osmotic balance^9,31,32^. Most notably for this work, intracellular potassium and potassium channels are well-established to be one of several mechanisms that regulate appropriate progression of the cell cycle^11,12^, and previous indications exist that potassium transport inhibition suppresses proliferation of cancer cells^33–36^.

We initially employed the picoeukaryote *Ostreococcus taur i*to test the hypothesis that circadian rhythms in potassium mechanistically couple the cell- and circadian cycles. This well-established minimal model cell for circadian rhythms^9,20^ shows the strongest coupling between the cell cycle and circadian cycle of all known eukaryotes^37^, making it ideal for studying the reciprocal interaction between potassium rhythms, the circadian TTFL, and cell proliferation. We then expanded our studies to further test the generality of the resulting hypotheses in mammalian cells, separated from *Ostreococcus* by ∼1 billion years of evolution^38^.

### Reciprocal feedback between potassium rhythms and clock gene expression

We first validated in *Ostreococcus* some key assumptions that underlie this investigation: 1) Intracellular potassium levels, [K^+^]_i_, oscillate in a circadian manner similarly to magnesium^9^ in the absence of environmental cues, peaking in the early subjective night. Other ions, such as calcium, remain constant over time (Fig. 1A); 2) potassium is highly abundant intracellularly with a large gradient over the plasma membrane (Extended Data Fig. 1A-B); 3) ion rhythms confer circadian rhythms in electrophysiological properties of the cell, as they do in mammalian cells^10^ (Extended Data Fig. 1C).

**Figure 1.**
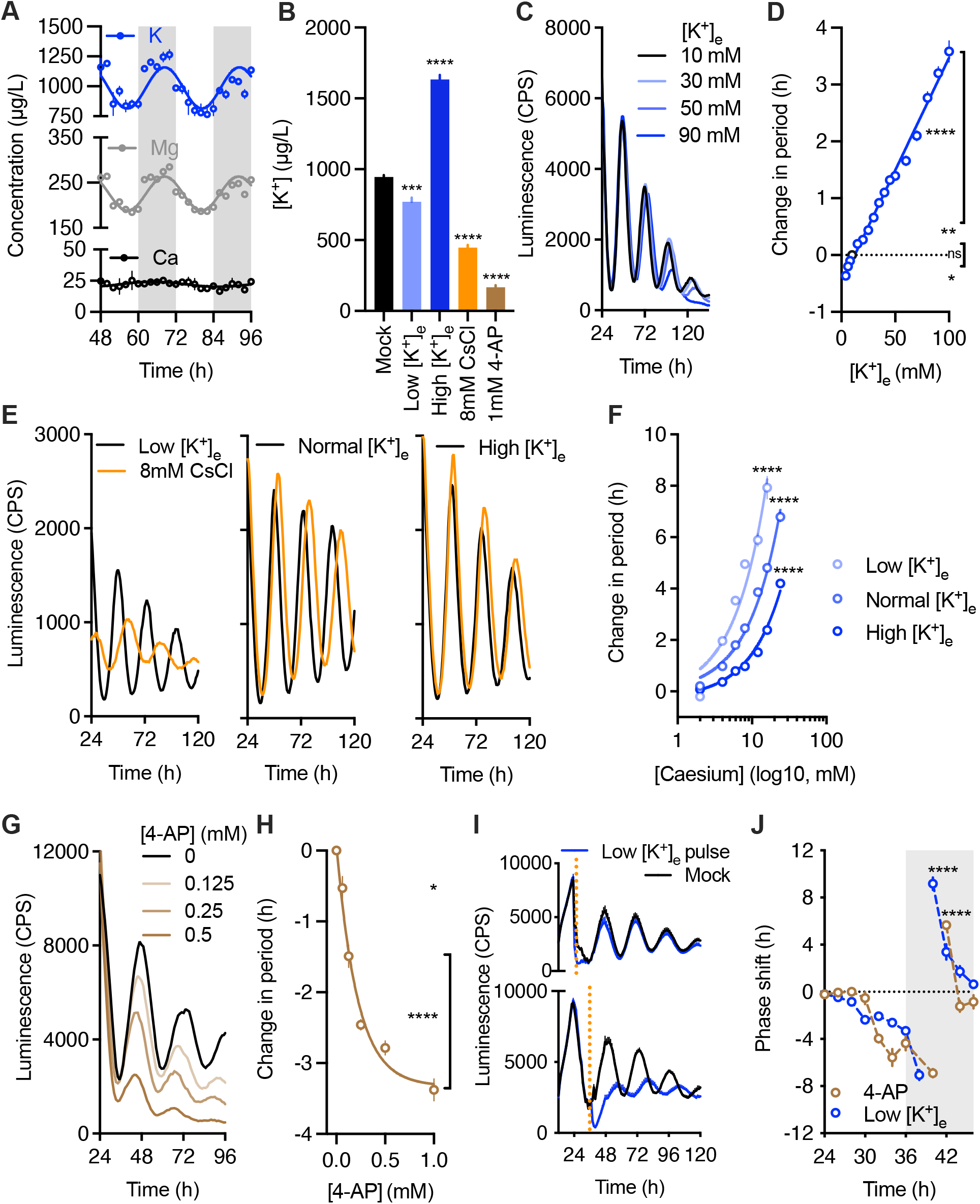
Potassium rhythms feed back to the circadian system. A) Quantification of intracellular. potassium, magnesium, and calcium concentrations in extracts taken over a time series under constant light conditions. Line represents a sine wave fitted through data points. n=4, mean±SEM. B) Changes in intracellular potassium levels can be induced by exogenous treatments. Samples collected after 16-18h of treatment. n=4, mean±SEM, one-way ANOVA, Dunnett’s multiple comparisons test vs. mock. C-D) Changes to circadian period quantified using the CCA1-LUC marker at different extracellular potassium concentrations, as example traces (C) or expressed as a dose response curve (D). n=16, mean±SEM, one-way ANOVA, Dunnett’s multiple comparisons test vs. 10 mM K^+^ control. E-F) Example traces (E) and dose response curve (F) for the effect of caesium on circadian period in media containing normal (10 mM), low (1 mM), or high (30 mM) extracellular potassium. n=8, mean±SEM, two-way ANOVA, caesium factor 72.2% of variation, Tukey’s multiple comparisons test. G-H) Example traces (G) and dose-response curve (H) for the effect of 4-AP on circadian period. n=8, mean±SEM, one-way ANOVA, Dunnett’s multiple comparisons test vs. mock. I) Examples of phase changes of the CCA1-LUC marker following low potassium pulses (orange dotted line) at 24h (subjective dawn, top) versus 36h (subjective dusk, bottom) under constant conditions. n=12. J) Phase response curves to pulse treatments with low extracellular potassium (blue) or saturating concentrations (1.5mM) of 4-AP (orange) in constant light. n=12, mean±SEM, one-way ANOVA, Tukey’s multiple comparisons test. P value summary only shown due to large number of comparisons. In A and J, grey area indicates subjective night. In D, F, H and J, period or phase changes are relative to control treatments.

Having satisfied these assumptions, to investigate reciprocal feedback between TTFL rhythmicity and changes in potassium levels, we established a set of treatments that affect [K^+^]_i_. Changing the concentration of extracellular potassium, [K^+^]_e_, induces corresponding changes in [K^+^]_i_ (Fig. 1B). [K^+^]_i_ can also be lowered using either caesium (Cs, a non-biological ion that competes with potassium for the same transport machinery^39^) or the voltage-dependent potassium ion channel inhibitor 4-aminopyridine (4-AP^40^). We then characterised the effect of these treatments on clock gene expression rhythms using an *in vivo* luminescence reporter of the TTFL clock gene, CCA1, fused to firefly luciferase (CCA1-LUC)^41^. Under constant light conditions (LL), increasing [K^+^]_e_ dose-dependently induced long period clock gene rhythmicity compared to the standard [K^+^]_e_ of 10 mM (Fig. 1C-D). Caesium dose-dependently induced period lengthening in media that contain the standard 10 mM [K^+^]_e_ (Fig. 1E-F). The effect of caesium can be modulated by changing extracellular potassium: a stronger effect is observed in low [K^+^]_e_ and a reduced effect in high [K^+^]_i_. This further confirms that period lengthening by caesium is indeed caused via direct competition with potassium. Unlike caesium, 4-AP induced short period clock gene rhythms (Fig. 1G-H). The differential effect on period observed between caesium and 4-AP treatments, both of which lower [K^+^]_i_ (Fig. 1B), is potentially due to their different modes of action: 4-AP acts specifically on voltage-gated potassium channels, while caesium competes with potassium non-selectively for transport and regulation of potassium-dependent intracellular processes^42^. Regardless, together these results reveal a direct and dose-dependent effect of potassium on the pace of the circadian TTFL.

To explore the time-dependency of the interaction between TTFL rhythms and potassium rhythms, we performed time series of 2h pulse treatments (Extended Data Fig. 1D) with low potassium under constant conditions. Interestingly, treatments at subjective dawn had no or modest effects on clock gene expression compared to controls, while treatments at subjective dusk produced a clear shift in phase (Fig. 1I). When circadian phase compared to mock treatments is analysed over a full 24 h cycle, the resulting Phase Response Curve (PRC) revealed a clear clock-gated response (Fig. 1J), with large phase shifts of up to 10 h following treatment during the early subjective night. This experiment was mirrored by 2 h pulsed treatment with a range of [4-AP], which confirmed the sensitive window during subjective night (Fig. 1J). The time of day where the greatest phase shifts are observed (Fig. 1J) coincides with the timing of peak potassium levels (Fig. 1A). These phase effects are not accompanied by large effects on circadian period (Extended Data Fig. 1E-F). Any period or phase effects described here or in the rest of this manuscript are not induced by the change in extracellular salinity: treatments used only induce minor changes in overall salinity (30-35 ppt) to which *Ostreococcus* clock gene expression is resilient (Extended Data Fig. 2A-B). Combined, these results indicate tight feedback between potassium and TTFL rhythms: potassium abundance is circadian-regulated and dose-dependently feeds back to regulate the period and phase of TTFL rhythmicity, fulfilling the definition of a regulator of cellular circadian rhythmicity.

### Specific intracellular potassium concentrations are required for optimal proliferation

The acute impact of reducing intracellular potassium on circadian phase during the subjective night (Fig. 1J) suggests that crucial mechanisms require the higher intracellular potassium levels found at this time (Fig. 1A). We therefore hypothesised that potassium rhythms might affect cell proliferation, as 1) cell cycle progression is tightly regulated and influenced by gradual changes in intracellular potassium^12^, and 2) strong coupling between the cell and circadian cycles is well-known across eukaryotes, although the mechanistic basis is poorly understood^7^. The *Ostreococcus* cell cycle is tightly gated by the circadian clock^37^ and the predominant phase of cell division is in the early night^8^. To assess the phase relationship between cell proliferation and potassium rhythms, we plotted approximate [K^+^]_i_ rhythms (from Fig. 1A) against cell cycle stages as inferred from proteome data^8^ and found that potassium built up from S phase (DNA-synthesis) through G2 (growth 2 phase) to peak around M phase (mitosis), and receded during G1 phase (Fig. 2A).

**Figure 2.**
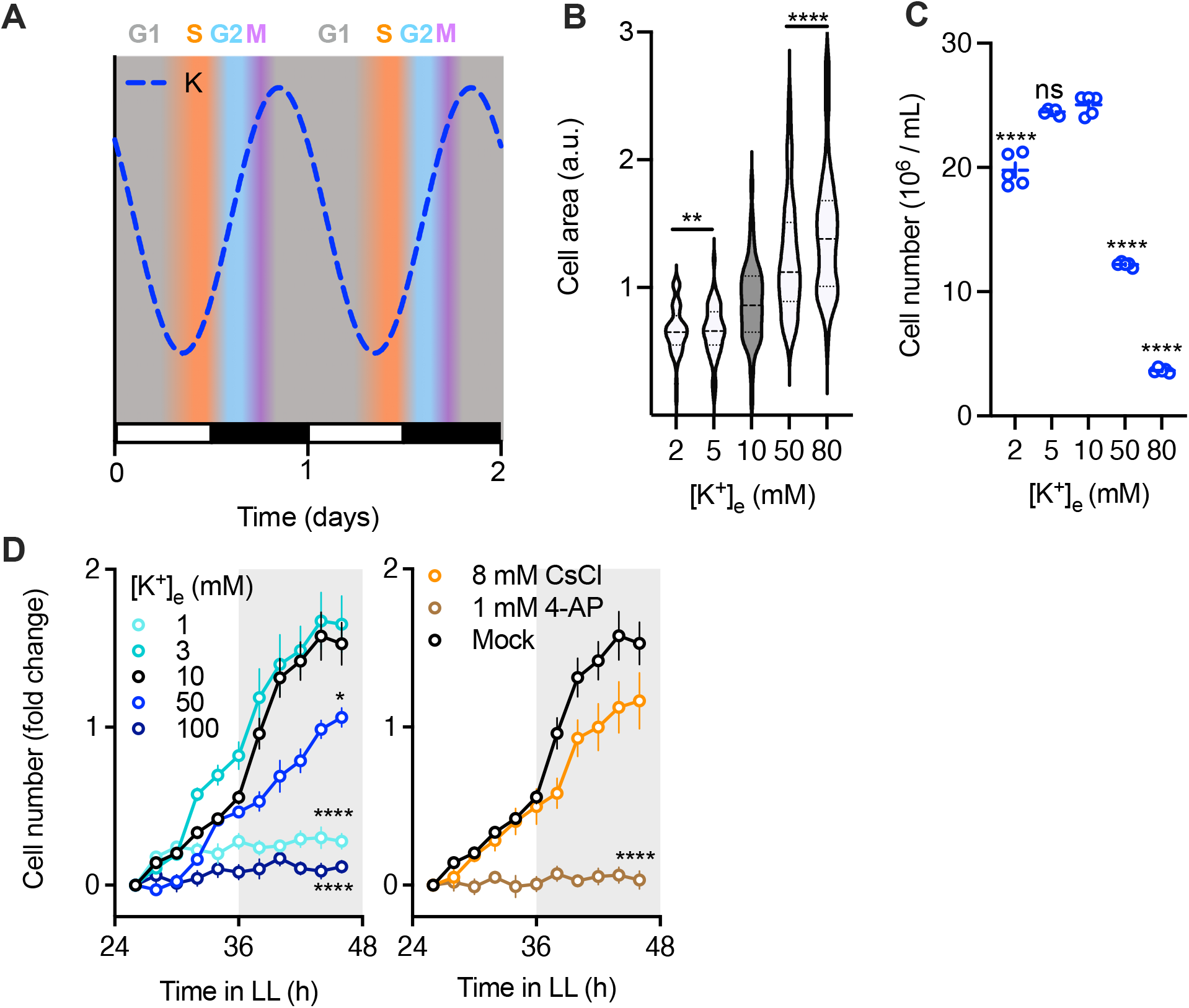
Cell proliferation is tuned by intracellular potassium concentration. A) Diagram indicating the phase correlation of rhythms in intracellular potassium concentration inferred from Figure 1A (blue dotted line), and the cell cycle phases inferred from proteome data^8^. Day and night phases are indicated by light and dark boxes. B-C) Cell size (B, n=50) or number (C, n=5) of cells grown for 5 days under differential potassium concentrations. Mean±SEM, one-way ANOVA, Dunnett’s multiple comparisons test vs. 10 mM K^+^ control. D) Cell proliferation of cells subjected to a range of potassium concentrations (left) or treatment with caesium or 4-AP (right), normalised to starting density. Grey areas represent subjective night periods. n=10, mean±SEM, Dunnett’s multiple comparison test vs 10mM K^+^ or mock control.

In cells like *Ostreococcus* and mammals, which lack a cell wall, increases in [K^+^]_i_ without compensatory decreases in other intracellular osmolytes^43^ will result in an associated increase in cell volume as water moves into the cell down the osmotic gradient. If [K^+^]_i_ would moderate intracellular osmolarity over the cell cycle, we predicted that cultures grown under a range of extracellular potassium concentrations would display differential cell volume and proliferation rates. Accordingly, high [K^+^]_e_ leads to unusually large cells and low [K^+^]_e_ to unusually small cells (Fig. 2B), both of which proliferate at a lower rate (Fig. 2C). Particularly dramatic effects are observed when increasing [K^+^]_e_ above the homeostatic [K^+^]_i_ (Extended data Fig. 1B), reversing the direction of the potassium gradient across the cell membrane. These effects are not due to changes in the total osmolarity of media but are specific to changes in potassium, as corresponding changes in sodium do not elicit a similar effect (Extended Data Fig. 2C). To further investigate the effect of potassium on the rate of cell proliferation, we monitored proliferation under treatments that affect [K^+^]_i_. Changing [K^+^]_e_ or addition of caesium or 4-AP led to a reduction or even a full arrest of cell proliferation (Fig. 2D). Again, the greater effect of 4-AP on proliferation is likely a result of its specific action at blocking K_v_1 channels, as opposed to the general competitive, but not inhibitory, action of caesium^44,45^. Taken together, our results suggest that cell proliferation and cell size in *Ostreococcus tauri* are highly sensitive to intracellular potassium levels and that rhythmic potassium levels within a tight concentration range are required for peak rates of cell division.

### Reciprocal interaction between the cell and circadian cycle

If cell proliferation is sensitive to a circadian rhythm in [K^+^]_i_ (Fig. 2), a possible hypothesis is that potassium rhythms mediate coupling between the cell and circadian cycles.

Having established the effects of modulating [K^+^]_i_ on clock gene expression (Fig. 1) and cell proliferation (Fig. 2), we next asked what effect modulating clock gene expression had on potassium and cell proliferation rhythms. Changing light fluence rates generally changes parameters of circadian TTFL rhythmicity^46,47^. We first tested the rhythmicity of the CCA1-LUC reporter under high light, low light, and darkness. We observed stable and high-amplitude gene expression rhythms under dim light (physiologically normal for *Ostreococcus*) and dampened and shorter period rhythms under high light (Fig. 3A-B). We saw no TTFL rhythmicity under complete darkness, due to the previously described inhibition of transcription under these conditions^48^. Rhythmicity of [K^+^]_i_ followed TTFL rhythmicity: oscillations occur at an advanced phase under dim compared to high light (Fig. 3C), while darkness induces apparent arrhythmia^9^. Cells subjected to different fluence rates also exhibited differential cell proliferation rates (Fig. 3D). While darkness (no clock gene expression, no potassium rhythms) led to a full cell cycle arrest as previously documented^37^, high light induced exceptionally high proliferation, indicating that cell proliferation can proceed unchecked when transcriptional rhythms are severely dampened. These results suggest reciprocal feedback between TTFL function and potassium rhythms, which combine to regulate cell proliferation.

**Figure 3.**
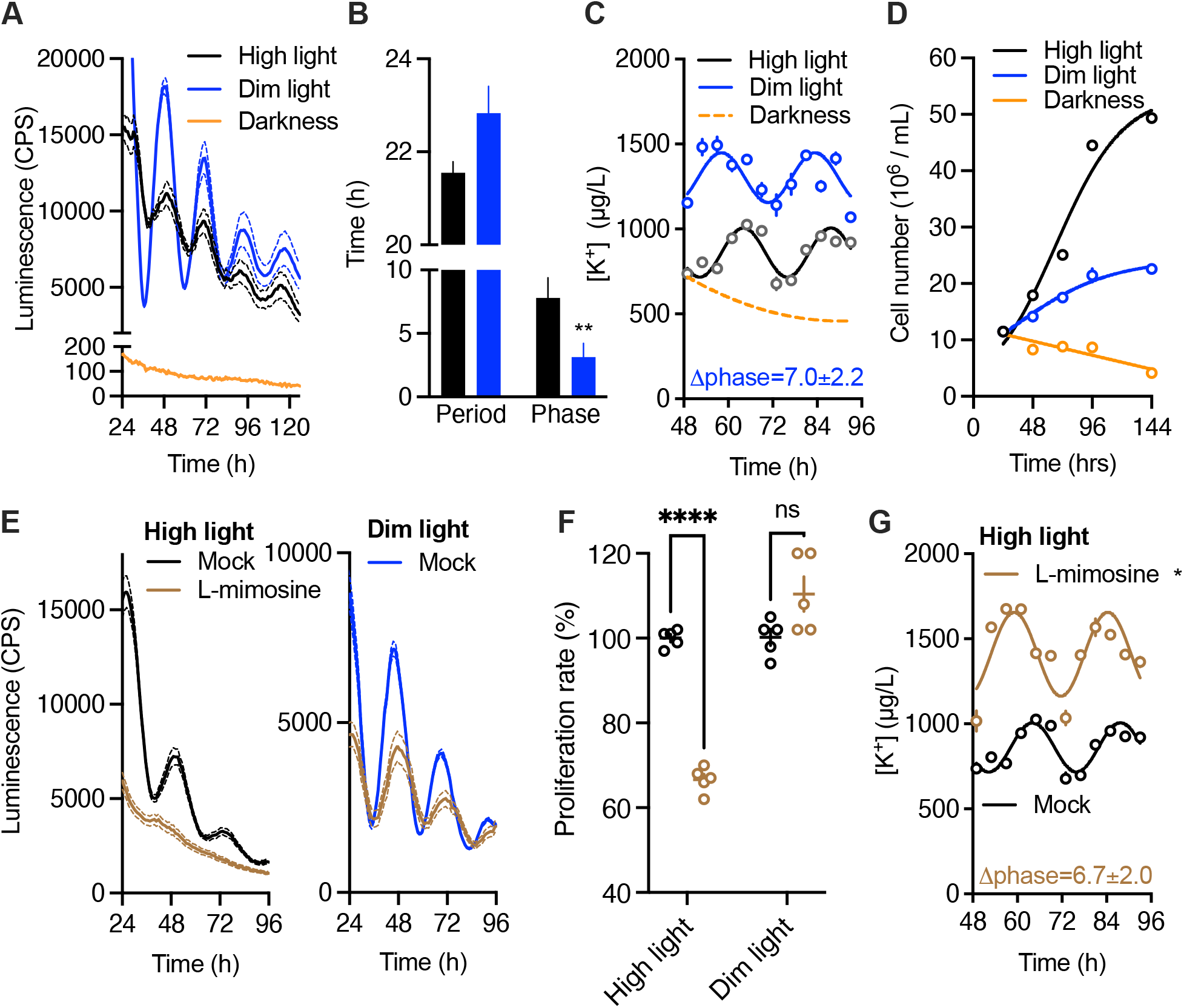
Reciprocal relationship between TTFL, potassium, and cell proliferation rhythms. A) Luminescence imaging of the clock reporter CCA1-LUC under different fluence rates. High light=16 µmoles/m^2^/sec; dim light=2 µmoles/m^2^/sec. n=10, mean±SEM. B) Quantification of period and phase of luminescence traces in (A); blue=dim light and black=high light. n=10, mean±SEM, two-way ANOVA. C) Intracellular potassium levels over time under high versus dim light conditions. n=3. Orange dotted line is a fit through previously published data from darkness^9^. Phase change provided for dim light vs. high light. D) Cell proliferation under different fluence rates. High and low light conditions fit to logistic growth curve, darkness preferentially fits to a straight line, extra sum-of-squares F-test, n=5. E) Luminescence of CCA1-LUC cultures under high (left panel) versus dim light (right panel), following a 24h treatment with 3.2 mM L-mimosine. n=14, mean±SEM. F) Cell proliferation of cultures subjected to the identical treatments as (E), counted one day after release of treatment. Unpaired t-test, Holm-Sidak, n=5, mean ±SEM. G) Potassium content of cultures subjected to L-mimosine versus controls under high light. Phase change provided for treated vs. mock conditions. High light control repeated from 3C. n=3, mean±SEM.

We then examined how inhibiting cell proliferation affected circadian gene expression and potassium rhythms. We used the non-protein amino acid inhibitor L-mimosine, which reversibly arrests cells at the G1/S interphase^49,50^. We asked whether inhibition had a differential effect under high versus dim light conditions. Following a 24h pulsed treatment (applied, then washed off), circadian gene expression was irretrievably lost under high light while it was only suppressed under dim light (Fig. 3E). Interestingly, cell proliferation was affected under high light but not dim light (Fig. 3F), indicating that inhibition of cell proliferation correlates with a lack of rhythmic gene expression. Inhibition leads to larger cell volumes under both light conditions but more strongly under high light (Extended Data Fig. 3A). As both TTFL rhythmicity and cell proliferation were inhibited by L-mimosine under high light, we then tested [K^+^]_i_ rhythms under those conditions. Remarkably, while [K^+^]_i_ oscillations occur at an advanced phase and elevated levelss, [K^+^]_i_ remains rhythmic under L-mimosine treatment (Fig. 3G). This indicates that potassium rhythms are independent of TTFL or cell cycle rhythmicity, as both these systems are arrhythmic under these conditions. This mirrors the earlier observation that potassium rhythms occur in human red blood cells^10^, which have no TTFL activity and do not divide.

### Enforced potassium gradients are sufficient to instruct cell proliferation

The results in Fig. 1-3 are consistent with potassium rhythms as a major link between circadian timing and the cell cycle. We therefore hypothesised that the effects of experimentally enforced changes in [K^+^]_i_ on cell proliferation should depend on circadian phase. In line with this, a 2h pulsed treatment (Extended Data Fig. 3B) of 4-AP applied at subjective dawn had no effect on cell proliferation, whereas it induced a large delay when applied at subjective dusk (Fig. 4A). Conversely, when pulsed treatments with high [K^+^]_e_ were given, treatment at subjective dusk had no effect on the subsequent proliferation pattern, while at dawn, a large phase advance in proliferation is observed after some initial cell death (Fig. 4B). Notably, treatments with high [K^+^]_e_ did not greatly affect TTFL rhythmicity at either time point (Extended Data Fig. 3C). A pulse of low [K^+^]_e_ did not greatly affect cell proliferation when applied at subjective dusk, while it induced cell death and an arrest of cell proliferation at subjective dawn (Fig. 4B). Interestingly, these strong effects on cell proliferation at subjective dawn occurs at the opposite circadian phase as that on TTFL rhythmicity (Fig.1I-J). These results highlight that changes in the potassium gradient are sufficient to advance or delay the timing of proliferation with respect to prior circadian phase.

**Figure 4.**
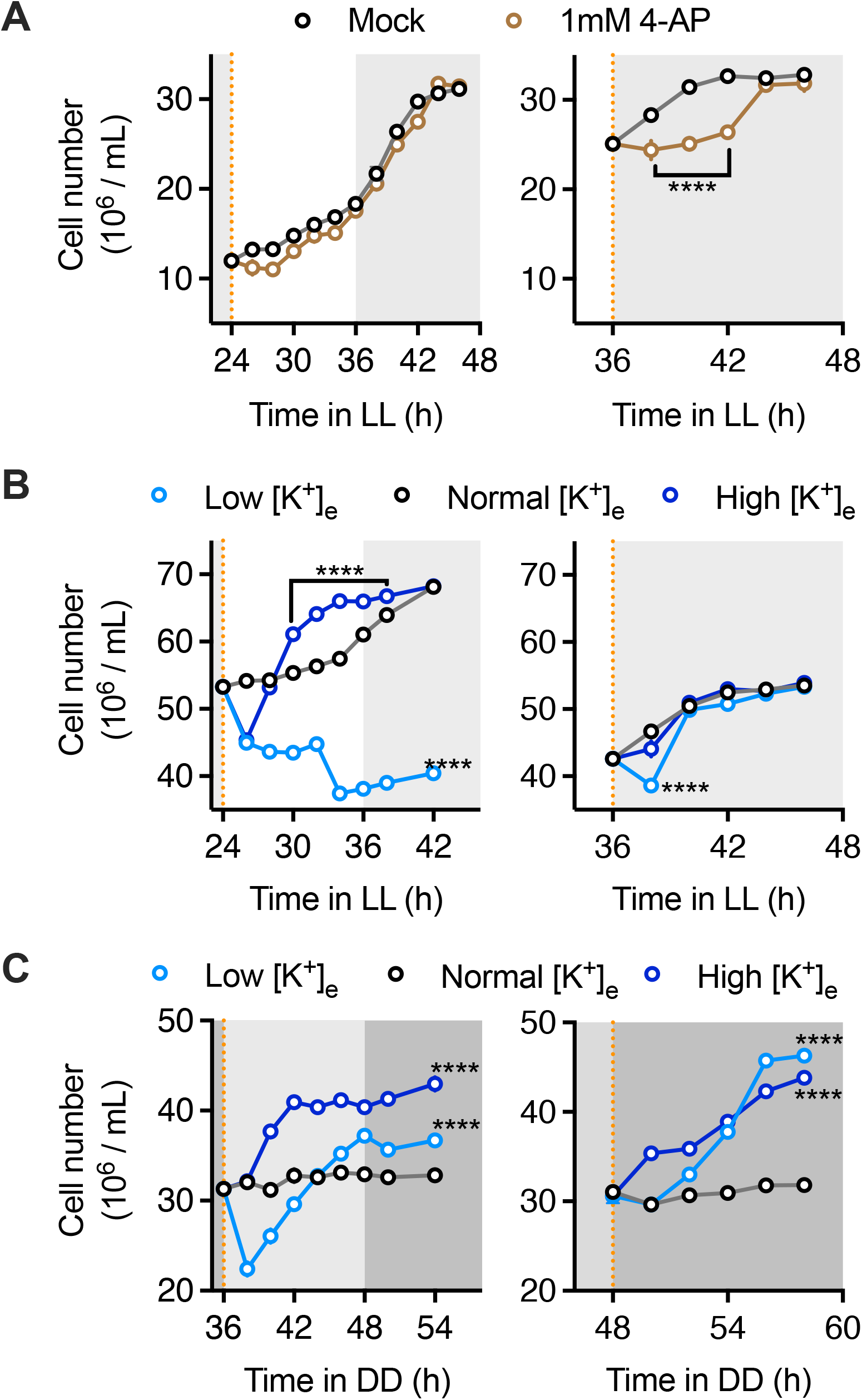
Potassium is sufficient to instruct cell division. A-B) Cell number upon 2h pulsed treatments with 4-AP (A) or low/high potassium (B) at subjective dawn (left panels) versus subjective dusk (right panels) under constant light conditions. C) Cell number upon 2h pulsed treatments with low/high potassium at subjective dawn (left panel) versus subjective dusk (right panel) under constant darkness. In all panels, darker areas represent subjective night and lighter areas subjective day; orange dotted lines indicate treatment times. n=6, mean±SEM, two-way ANOVA, Dunnett’s multiple comparisons test vs. control. Significance markers in middle of line or brackets indicate regions of a line that differ from control, significance markers at the end of a line indicate that all points on a line differ from control.

As potassium gradients affect cell proliferation in a manner that does not necessarily relate to TTFL rhythmicity, we then asked whether potassium could also induce changes in the cell cycle under complete darkness, when gene expression, potassium, and proliferation are all arrhythmic (Fig. 3A-D). Pulse treatments with either high or low potassium do not restore TTFL rhythms under these conditions (Extended Data Fig. 3D). However, a rapid ∼50% increase of the total cell number was induced by treatments either at subjective dawn or dusk (Fig. 4C). The effects in Fig. 4B-C become more obvious when plotted relative to control treatments and correcting for initial cell death (Extended Data Fig. 4E). These results establish that 1) a sudden, enforced potassium gradient is sufficient to instruct at least one round of cell division and 2) that a functional TTFL is required to deliver phase sensitivity (i.e. to correctly translate a change in potassium into committing to cell proliferation).

### Potassium affects timekeeping in mammalian cells

To test the generality of our findings, we asked how closely the results from model *Ostreococcus* cells (Fig. 1-4) correspond to mammalian systems. Previous work has identified coupling between the cell cycle and circadian rhythms in vertebrates^24,25,51^, including direct circadian regulation of cell cycle regulators such as Wee1 kinase and cyclins^25,52^. Reciprocally, cell cycle regulators, including the master tumour suppressor p53, actively regulate mammalian circadian rhythmicity by repressing expression of the critical clock gene Period (PER)^53^.

First, the effects of treatment with 4-AP and caesium on the mammalian TTFL system was assessed in cells expressing circadian bioluminescence reporters. In actively dividing NIH 3T3 cells expressing luciferase under the circadian *Per2* promoter (Per2:LUC), manipulation of intracellular potassium dose-dependently lengthened circadian gene expression (Fig. 5A-D, Extended Data Fig. 4A-B). To confirm relevance of our findings in primary cells, we repeated these experiments in adult lung fibroblasts derived from the PERIOD2-LUCIFERASE (PER2-LUC) mouse, where luciferase is expressed in fusion with the endogenous PER2 protein^54^. In both actively dividing (Extended Data Fig. 4C-F) and confluent (Extended Data Fig. 4G-L) cells, we observed changes in period comparable to both NIH 3T3 cells as well as *Ostreococcus* cells. Next, we tested whether potassium affects clock gene expression in a circadian phase dependent manner. Treatments that affect potassium were applied at the peak versus the trough of PER2 expression in fibroblasts as [K^+^]_i_ peaks a few hours before PER2 in these cells^26^. Caesium treatment at the peak of PER2 expression (i.e. decreasing [K^+^]_i_ phase) did not affect phase, while treatments at the trough of PER2 (increasing [K^+^]_i_ phase) clearly shifted circadian phase compared to controls (Extended Data Fig. 4M-N). This compares favourably to *Ostreococcus*, which also showed peak sensitivity to potassium perturbation at the time of peak [K^+^]_i_ (Fig. 1J). In contrast, 4-AP induced phase shifts at both timepoints (Extended Data Fig. 4O-P), unlike in *Ostreococcus* (Fig. 1J). We speculate this is likely due to differences in relative expression and utilisation of voltage-gated potassium channels between the two organisms. Together, these results indicate that circadian rhythms in potassium levels in mammalian cells, as in *Ostreococcus,* feed back to regulate TTFL gene expression in a phase-dependent manner.

**Figure 5.**
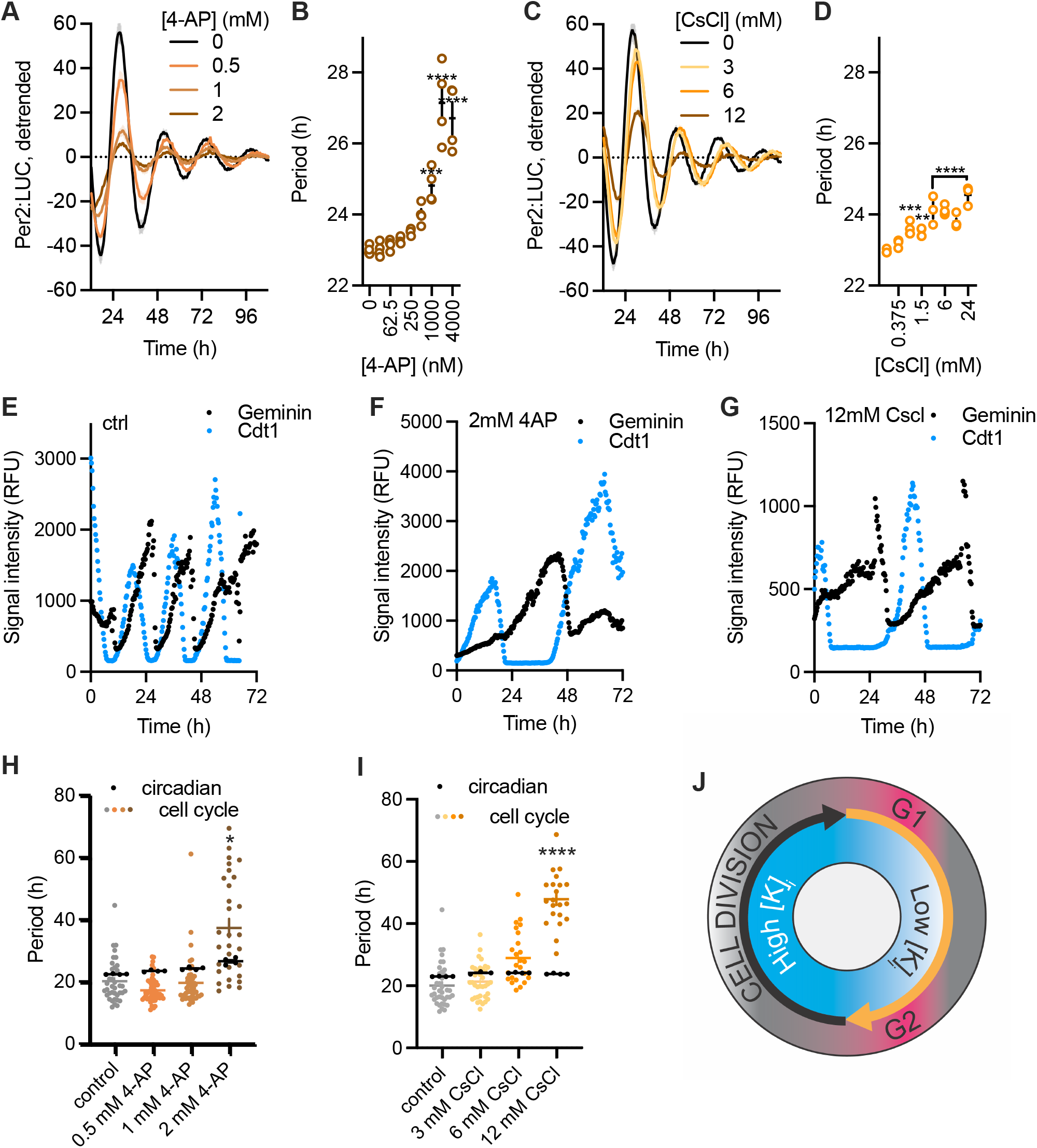
Potassium regulates both cell and circadian cycles in mammalian cells. A-D) Actively dividing Per2:LUC NIH 3T3s treated with increasing concentrations of 4AP (A-B) or CsCl (C-D) show increasing circadian period. n=3, mean±SEM, one-way ANOVA, Dunnett’s multiple comparisons test vs. control. E-G) Representative traces of FUCCI markers from NIH 3T3s under control conditions (E) or treated with 2 mM 4AP (F) or 12 mM CsCl (G). H-I) Comparison of the change in period of the circadian marker and cell cycle marker under increasing concentrations of 4-AP (H) or CsCl (I). n≥21 nuclei taken from at least 3 independent wells, mean±SEM. Fisher’s LSD test. J) A general model for the relationship between potassium rhythms and cell division.

### Potassium affects cell proliferation in mammalian cells

To determine whether manipulation of potassium also inhibits cell proliferation in mammalian cells, PER2-LUC fibroblasts were subjected to treatment with 4-AP or caesium at peak versus trough PER2 levels. Although cell proliferation is inhibited by potassium manipulations at either phase, a significantly greater effect is observed upon treatments at peak PER2 levels (Extended Data Fig. 5A-F). Notably, as in *Ostreococcus,* the maximal effects of these treatments on TTFL rhythmicity and on cell proliferation occur at opposite circadian phases. Overall, the results in Fig. 5A-D and Extended Data Fig. 4 and 5A-F establish that modulating potassium affects TTFL rhythmicity as well as cell proliferation in mammalian cells in a phase-dependent manner.

Although much weaker than that observed in *Ostreococcus,* 1:1 circadian:cell cycle coupling, such that there is one cell division per circadian cycle, was previously demonstrated in freely proliferating mammalian cells under standard culture conditions in 10% serum^24,25^. We posited that if intracellular potassium was acting to couple these two cyclical processes, then manipulation of potassium should disrupt this coupling such that the two processes cease to run in parallel. To test this, we employed NIH 3T3 cells expressing the FUCCI-2A system, which allows for non-invasive imaging of cell cycle progression through expression of fluorescently-tagged fragments of Cdt1 (accumulating in G1) and Geminin (accumulating in S/G2/M)^55^. Single cell imaging (Supplemental Data Files 1-3) confirmed a mean period of cell division (Fig. 5E) that aligns with the circadian period of these cells (Fig. 5B, D). This conforms to previous work^25^ that demonstrated coupling between the circadian and cell cycles in these cells under these conditions. Cells treated with 4-AP or CsCl show slower cell cycle rhythms (Fig. 5F, G), a decreased rate of proliferation (Extended Data Fig. 5G-J) and an increased proclivity towards cell cycle arrest (Extended Data Fig. 5K, L). When subjected to a range of concentrations of 4-AP or CsCl, a strong dose-dependent increase in the average cell cycle period is observed (Fig. 5H-I, coloured data points). Importantly, these increases in cell cycle period were much stronger than, and did not correlate with, the observed increase in circadian period at the same concentrations of these drugs (black data points). These results are therefore consistent with our prediction that disrupting potassium transport uncouples the circadian clock from cell cycle progression. Taken together, our results strongly suggest that loss of circadian-cell cycle coupling is a general consequence of disruption of intracellular potassium levels in mammalian cells.

## Discussion

Previous studies have largely focused on links between cell division and the transcriptionally driven circadian TTFL. However, increasing knowledge of robust and universal circadian rhythms that continue under non-transcriptional conditions^9,19,20,56^ demonstrate that there is a need to shift the focus to post-translational events to clearly unravel the potential link(s) between the circadian and cell cycles. Oscillatory intracellular potassium is one of these cellular properties that appear to run in parallel with TTFL rhythms. This work demonstrates that intracellular potassium levels regulate circadian rhythms and coordinate phase-dependent cell cycle progression even in the absence of transcriptional timekeeping. Furthermore, these results establish that the evolutionarily conserved circadian rhythms in potassium levels^9,26^ are a critical factor coupling the cell and circadian cycles (Fig. 5J).

Our previous work showed that circadian rhythms in intracellular ion concentration buffer intracellular osmolarity in response to daily rhythms in protein synthesis and abundance in confluent cells^26^. Considering this, it is tempting to postulate that in proliferating cells, similar mechanisms are required to regulate cell volume changes in response to protein synthesis during G1 phase^57,58^. In line with this, G1 phase sees a dramatic reduction in [K^+^]_i_ (Fig. 2A, 5J) when protein synthesis rates are highest. The increase in potassium during G2 phase, and the consequent increase in intracellular osmolarity, is likely to contribute to the ingress of water required for cell growth ahead of cytokinesis.

Interestingly, abnormal proliferation phenotypes and cell properties found in multiple cancer cells including glioma, hepatoblastoma, breast cancer or malignant astrocytoma are correlated with altered K^+^ homeostasis, or with the altered polarization profiles that these cause^33–36,59–61^. The inhibition of K^+^ transporters in tumour cells also showed a strong therapeutic potential in cancer treatments, highlighting this as a potential therapy target for cancer research^59,62^. It is also worth noting that disruption of circadian rhythms is permissive for the development of cancer^16^. The novel fundamental insights reported here into the integration of potassium rhythms, TTFL rhythms, and cell proliferation can inform research into the underlying molecular regulation of these coupled systems, ultimately contributing to future cancer research and therapy.

## Acknowledgements

The authors thank Erin Henslee and Fatima Labeed for expertise and equipment required for DEP measurements in Extended Data Figure 1C. We also thank Seth Rubin for his critical review of the manuscript, Franck Delaunay for his kind gift of Rev-Erbα-Venus FUCCI-2A NIH 3T3 cells, and David Welsh for the generous donation of PERIOD2-LUCIFERASE mouse lung tissue.

## Funding

SGR and GvO were supported by a Wellcome Trust Discovery Award (225212/Z/22/Z) and a Leverhulme Trust Research Grant (RPG-2019-184). LLH and EG were supported by Wellcome Trust Institutional Strategic Support Fund awards to GvO. PC and CLP are supported by US NIH grant R35 GM141849. The UCSC Chemical Screening Center (RRID SCR_021114) was supported by a National Institutes of Health High End Instrumentation Grant (1S10OD028730-01A1). AS is funded by the Deutsche Forschungsgemeinschaft (DFG) under Germany’s Excellence Strategy EXC 2030 (390661388) and a project grant (510582209).

## Methods

### Ostreococcus tauri methods

#### Bioluminescence recording

Transgenic CCA1-LUC *Ostreococcus* cells^41^ were grown in vented tissue culture flasks (Sarstedt) at 20°C in artificial sea water (ASW) as described previously^9^ with a light intensity of 17 μmol m^-2^ s^-1^ under blue light filters (183 Moonlight Blue Filter, Lee filters). Cells were entrained under 12h light/12h dark (LD) cycles for 6-7 days to reach optimal cell density for experimental use. CCA1-LUC cell cultures were diluted 1:3 in fresh ASW 30-35 ppt and supplemented with 0.2 mM D-luciferin. 90 µL was added to wells of a 384-well plate (Greiner) and imaged in a luminescence plate reader (Berthold TriStar2) under 2 μmol m^-2^ s^-1^ blue light (183 Moonlight Blue Filter, Lee filters) for 5-7 days under constant light conditions. 4-AP (Sigma) or CsCl (Sigma) treatments were added at 10x concentration. For washout treatments, media from each well was carefully removed and replaced with media containing the appropriate treatment and luciferin, avoiding disturbing the cell aggregates. After the pulse treatments, cells were fully resuspended with fresh ASW media + luciferin. Treatments were performed on the second day of constant conditions (24h into LL or 36h into DD) and all fall within a range of overall salinity of 30-35 ppt. Salinity experiments were performed by adjusting [NaCl]. For differential light conditions, light intensity was adjusted at the start of imaging. High light = 17 μmol m^-2^ s^-1^ and dim light = 2 μmol m^-2^ s^-1^. Results were analysed and plotted using GraphPad Prism v9. Period and phase analyses were performed using BioDare2^63^ as previously published^8^.

### Cell number and area analysis

For cell number and cell area analyses, *Ostreococcus* cells subjected to identical experimental settings and conditions as for the luminescence assays were harvested and counted using a haemocytometer with a light microscope on 40x magnification. Data was collected from at least 5 technical replicates for each time point. For cell area, cell samples were quickly mixed 1:1 with −80°C methanol (70%) to preserve cell volume. Pictures were taken with a light microscope from at least 100 cells for each condition and cell area was measured with ImageJ.

### Ion analysis

For ion analyses, 25-30 ml *Ostreococcus* cell cultures were collected at stated times, pelleted, and washed twice with 1 M Sorbitol (Sigma) to remove all salts present in the media. Pellets were then resuspended in 100 µL of 69% Nitric acid (MERCK) and digested O/N at RT. Samples were diluted to 5% Nitric acid with HPLC grade water. 4 technical replicates for each time point were analysed using Microwave Plasma - Atomic Emission Spectrometry (MP-AES 4210, Agilent) as reported previously^24^.

### Dielectrophoresis

For dielectrophoresis (DEP), *Ostreococcus* cultures were entrained under 14h light:10h dark cycles before transfer into constant conditions. Starting at 24h, every 4 hours, 15 mL of 27.5 × 10^6^ cells/mL cultures were transferred to 15 mL falcon tube and washed twice in 15 mL iso-osmotic 1 M sorbitol by centrifuging for 2 minutes at 4472 *g*, and then resuspended in a final volume of 0.5 mL 1 M sorbitol. 75 µL of these cell suspensions were pipetted into 3DEP chips (DEPtech, Heathfield, UK), which were subsequently inserted into a 3DEP reader (DEPtech). Pin connections energised each well at 10 Vp–p, with a different frequency applied to each of the 20 wells and with the wells collectively energised for 10 s at five points per decade (10 kHz—20 MHz). This was repeated at each time point at least three times. The raw data were fitted with a single-shell model in order to extract the electrophysiological parameters as previously^10,64^, accepting spectra producing R^2^ values of 0.9 or greater.

### Mammalian cell methods Isolation of cell lines

NIH 3T3 Rev-Erbα-Venus FUCCI-2A cells were a gift from Franck Delaunay. NIH 3T3 Per2:LUC cells were generated previously^65^. PERIOD2-LUCIFERASE^54^ lung fibroblasts were derived from mouse lung kindly donated by David Welsh. Mouse primary fibroblasts were isolated according to an established protocol^66^. For this, lung tissue was stored in ice-cold PBS for up to 24 hrs. Tissue samples were subsequently removed from PBS and cut in to ∼1 mm^3^ sections using a pair of sterile scalpels, before being transferred to a 50 mL falcon tube with 10 mL “digestion medium” (DMEM/F12 supplemented with pen/strep, Mycozap Plus PR and 0.14 U/mL Liberase) and incubated at 37°C, stirring slowly, for 30 min, or until the tissue fragments turned white. The tissue fragments were then titurated using a 10 mL pipette and 40 mL “initial culture medium” (DMEM/F12, supplemented with pen/strep, Mycozap Plus PR and 15% HyClone FetalClone III) added before the tube was centrifuged at 700x g for 5 min. The resulting supernatant was discarded, the pellet resuspended in a further 20 mL “initial culture media” and the tube centrifuged for a further 5 min. The supernatant was again discarded, the pellet resuspended in 10mL “initial culture media” and transferred to a 10 cm tissue culture dish and incubated at 37°C, 5% CO_2_, 3% O_2_. After 7 days, media was refreshed and after a further 7 days, cells were split and re-plated in “selection medium” (MEM supplemented with pen/strep, non-essential amino acids, sodium pyruvate and 10% HyClone FetalClone III). After a further 2 weeks, cells were transferred to DMEM-based culture medium (DMEM supplemented with pen/strep and 10% HyClone FetalClone III). Immortalization was achieved by serial passage of cells at 37°C, 5% CO_2_, 20% O_2_. Cell lines were authenticated by observation of morphology and by continued expression of the bioluminescent reporter.

### Bioluminescence recording

For bioluminescence assays, cells were grown to confluence in 12 well or 35mm dishes in high-glucose (27.8 mM), glutamax-containing DMEM (GIBCO) supplemented with 10% serum (HyClone FetalClone III, Themofisher) and pen/strep and subjected to temperature cycles of 12 hours 37°C followed by 12 hours at 32°C. Confluent cultures were kept for up to 4 weeks with media refreshed every 7-10 days. For recording, cells were changed to MOPS-buffered “Air Media” (Bicarbonate-free DMEM, 5mg/mL glucose, 0.35 mg/mL sodium bicarbonate, 0.02 M MOPS, 100 μg/mL pen/strep, 1% Glutamax, 1 mM luciferin, pH 7.4, 325 mOsm)^67^. 10% serum was used in all cases except for low [K]_e_ experiments. Cells were then transferred to a Lumicyle (Actimetrics) or an ALLIGATOR (Cairn Research), where bioluminescent activity was recorded at 15 min intervals using an electron multiplying charge-coupled device (EM-CCD) at constant 37°C. For treatment, cells were removed from recording on a heatpad and kept at constant 37°C for treatment before returning to recording. Detrending of bioluminescent traces, where appropriate, was performed using a 24 h moving average detrend. Bioluminescent traces from mammalian cells were fitted with damped cosine waves in Prism 10 (GraphPad) using the following equation:

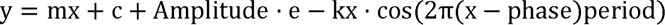

where y is the signal, x the corresponding time, amplitude is the height of the peak of the waveform above the trend line, k is the decay constant (such that 1/k is the half-life), phase is the shift relative to a cos wave and the period is the time taken for a complete cycle to occur.

### Fluorescence imaging and analysis

For fluorescence cell imaging of Rev-Erbα-Venus FUCCI-2A NIH 3T3s, cells were plated to ∼7% confluency in a black, clear-bottomed 96-well plate in MOPS-buffered ‘Air media’ with 10% serum but lacking luciferin (see above) with the indicated concentrations of CsCl, NaCl or 4AP. Cells were then maintained at constant 37°C for 36 hours before moving to an Opera Phenix Plus (Revvity Perkin Elmer, at the UCSC Chemical Screening Center RRID SCR_021114)) for recording at constant 37°C in a humidified, dark environment. Cells were imaged every 18 minutes with 9 fields of view per well, using a 10x air objective, two peak focusing at the −5 um focal plane, 50 um pinhole spinning disc, 2160×2160 px camera, binning 2, and the following channels: brightfield with transmitted light and a 650-760 nm filter with 20 ms exposure at 20% light power; MKO2 with a 561 nm excitation laser and a 571-596 nm emission filter with 60 ms exposure at 95% laser power; and E2-Crimson with a 640 nm laser and a 650-760 nm emission filter with 800 ms exposure at 95% laser power.

For fluorescence imaging of PER2-LUC fibroblasts, cells were plated to ∼7% confluency in a black, clear-bottomed 96 well plate in high-glucose (27.8 mM), glutamax-containing DMEM (GIBCO) supplemented with 10% serum (HyClone FetalClone III, Themofisher) and pen/strep and maintained under temperature cycles as described above. After 24 hours, they were treated with Cell Tracker Red CMTPX Dye (Thermo Fisher) for 30 minutes before changing to MOPS-buffered ‘Air medium’ with 10% serum but lacking luciferin and moving to an Opera Phenix Plus (Revvity Perkin Elmer, at the UCSC Chemical Screening Center RRID SCR_021114)) for recording at constant 37°C in a humidified environment. Cells were imaged every 15 minutes using the same hardware as described above for Rev-Erbα-Venus FUCCI-2A 3T3s, except the focal plane was at −8 um and the following channels: brightfield with transmitted light and a 650-760 nm filter with 100 ms exposure at 40% light power; Cell Tracker Red with a 561 nm excitation laser and a 570-630 nm emission filter with 100 ms exposure at 95% laser power.

Fluorescence image analysis of 3T3s and primary fibroblasts was performed using the cell tracking feature of Harmony 5.1 high-content imaging and analysis software (Revvity Perkin Elmer, at the UCSC Chemical Screening Center (RRID SCR_021114)). For Rev-Erbα-Venus FUCCI-2A 3T3s, the images were flatfield corrected, the sum of the three fluorescent channels calculated, and a sliding parabola applied, from which nuclei were identified, nuclei intersecting with the image border removed, the nuclei then tracked (with tracked nuclei requiring an overlap of at least 1% between sequential images), and the intensities of the three fluorescent channels measured for each nucleus. For subsequent analysis, only cells with continuous tracks longer than 200 images were used. Senescent cells were defined as those cells that did not undergo a cell division event during the recording, determined as those cells which did not complete a cycle of cdt1 and geminin expression from the FUCCI-2A reporter. Period of cell division was determined from those cells that underwent at least two full cycles of cell-cycle gene expression, with period quantification performed using BioDare 2^60^.

For Cell Tracker Red images, PER2-LUC fibroblasts were tracked as described above for the 3T3s except the Cell Tracker Red raw image was used for cell identification, and the number of cells was outputted.

### Statistics

All statistical analysis was performed using Prism 10 (GraphPad).

**Extended Data Figure 1. The *Ostreococcus* ionome**

A) MP-AES quantification of intracellular ions at subjective day versus subjective night in constant light conditions. n=3, mean±SEM, two-way ANOVA with Holm-Sidak’s multiple comparisons. B) Potassium concentrations in extracts were converted to intracellular concentration by correcting dilution rate, total cell number in each sample, and the known average cell volume^68^. Known concentrations in artificial sea water are provided. C) Dielectrophoresis results show rhythmicity in the electophysiological properties of the cell (σ_cyt_ = cytoplasmic conductivity; G_eff_ = effective membrane conductance). D) Schematic representation of treatments in main Figure 1I-J. E-F) Period analyses of the phase response curves in Fig. 1J (E: low potassium, F: 4-AP). n=12, box and whiskers; min to max.

**Extended Data Figure 2. *Ostreococcus* is insensitive to moderate changes in extracellular salinity**

A) Period analyses of luminescence traces under constant light conditions of CCA1-LUC cells subjected to media with differential salinity. n=8, mean±SEM, one-way ANOVA with pairwise comparisons to control conditions (30ppt). B) Cell counting of cells subjected to with differential salinity. n=4, mean±SEM, one way ANOVA with pairwise comparisons to control conditions (30ppt) for endpoint only. For A-B: potassium treatments performed throughout this manuscript do not exceed the salinity range of 30-35ppt (indicated by orange shading), showing no significant changes in circadian rhythms or cell division.

**Extended Data Figure 3. Additional treatments of *Ostreococcus* cells**

A) Light microscopy pictures (left; each panel is composed of six separate pictures) and quantification of cell size changes (right) upon treatment with 3.2mM L-mimosine in high versus dim light. n=15, mean±SEM, one-way ANOVA with pairwise comparisons. B) Diagram depicting the experimental designs of treatment pulses in Fig. 4. C) Luminescent imaging of CCA1-LUC cells subjected to 2h pulses of high potassium under constant light at subjective dawn (left panel) versus dusk (right panel). Treatment time indicated by orange dotted line. n=16, mean±SEM. D) Luminescent imaging of CCA1-LUC cells subjected to 2h pulses of low/high potassium under darkness. n=8, mean±SEM. E) Changes in cell number under light (functional TTFL) versus darkness (no TTFL) in response to pulsed treatments with low (bottom) or high potassium (top); identical data as in main Figure 4B and C, but expressed relative to the control mock treatments and corrected for initial cell death.

**Extended Data Figure 4. Effect of potassium manipulation on mammalian circadian period and phase**

A) Actively dividing Per2:LUC NIH 3T3s treated with increasing concentrations of 4-AP show increasing circadian period. n=3, mean±SEM. Extended from Fig. 5A. B) Actively dividing Per2:LUC NIH 3T3s treated with increasing concentrations of CsCl show increasing circadian period. n=3, mean±SEM. Extended from Fig. 5C. C-D) Actively dividing PER2-LUC fibroblasts treated with increasing concentrations of 4-AP show increasing circadian period. n=3, mean±SEM, asymmetric dose-response curve. E-F) Actively dividing PER2-LUC fibroblasts treated with increasing concentrations of CsCl show increasing circadian period. n=3, mean±SEM, asymmetric dose-response curve. G-H) Confluent PER2-LUC fibroblasts treated with increasing concentrations of 4-AP show increasing circadian period. n=3, mean±SEM, asymmetric dose-response curve. I-J) Confluent PER2-LUC fibroblasts treated with increasing concentrations of CsCl show increasing circadian period. n=3, mean±SEM, asymmetric dose-response curve. K-L) Confluent PER2-LUC fibroblasts treated with increasing concentrations of extracellular potassium show increasing circadian period. n=3, mean±SEM, asymmetric dose-response curve. M) Treatment with 6 mM CsCl induces a shift in circadian phase that varies with phase of application. n=3, mean±SEM. N) Quantification of phase shift versus NaCl control, t-test. O) Treatment with 1 mM 4-AP induces a shift in circadian phase that varies with phase of application. n=3, mean±SEM. P) Quantification of phase shift versus vehicle control, t-test.

**Extended Data Figure 5. Effect of potassium manipulation on mammalian cell proliferation**

A-C) Treatment of actively dividing PER2-LUC fibroblasts with 1 mM 4-AP at different phases results in differing effects on subsequent log-phase proliferation rate. n≥4, mean±SEM, Šidák’s multiple comparison’s test. D-F) Treatment of actively dividing PER2-LUC fibroblasts with 6 mM CsCl at different phases results in differing effects on subsequent log-phase proliferation rate. n≥4, mean±SEM, Šidák’s multiple comparison’s test. G-H) Treatment of phase-unsynchronised FUCCI-2A NIH 3T3s with 4-AP reduces proliferation rate in a dose-dependent manner. n≥4, mean±SEM, Dunnett’s multiple comparisons test. I-J) Treatment of phase-unsynchronised FUCCI-2A NIH 3T3s with 4-AP reduces proliferation rate in a dose-dependent manner. n≥4, mean±SEM, Dunnett’s multiple comparisons test. K-L) Percentage of cells that failed to complete a division event under increasing concentrations of 4-AP (K) or CsCl (L). n=3, mean±SEM, Dunnett’s multiple comparisons test.

## Notes

### Competing Interest Statement

The authors have declared no competing interest.

