## Supplementary figures and images for "Potassium rhythms couple the circadian clock to the cell cycle"

### Extended Data Figure 1

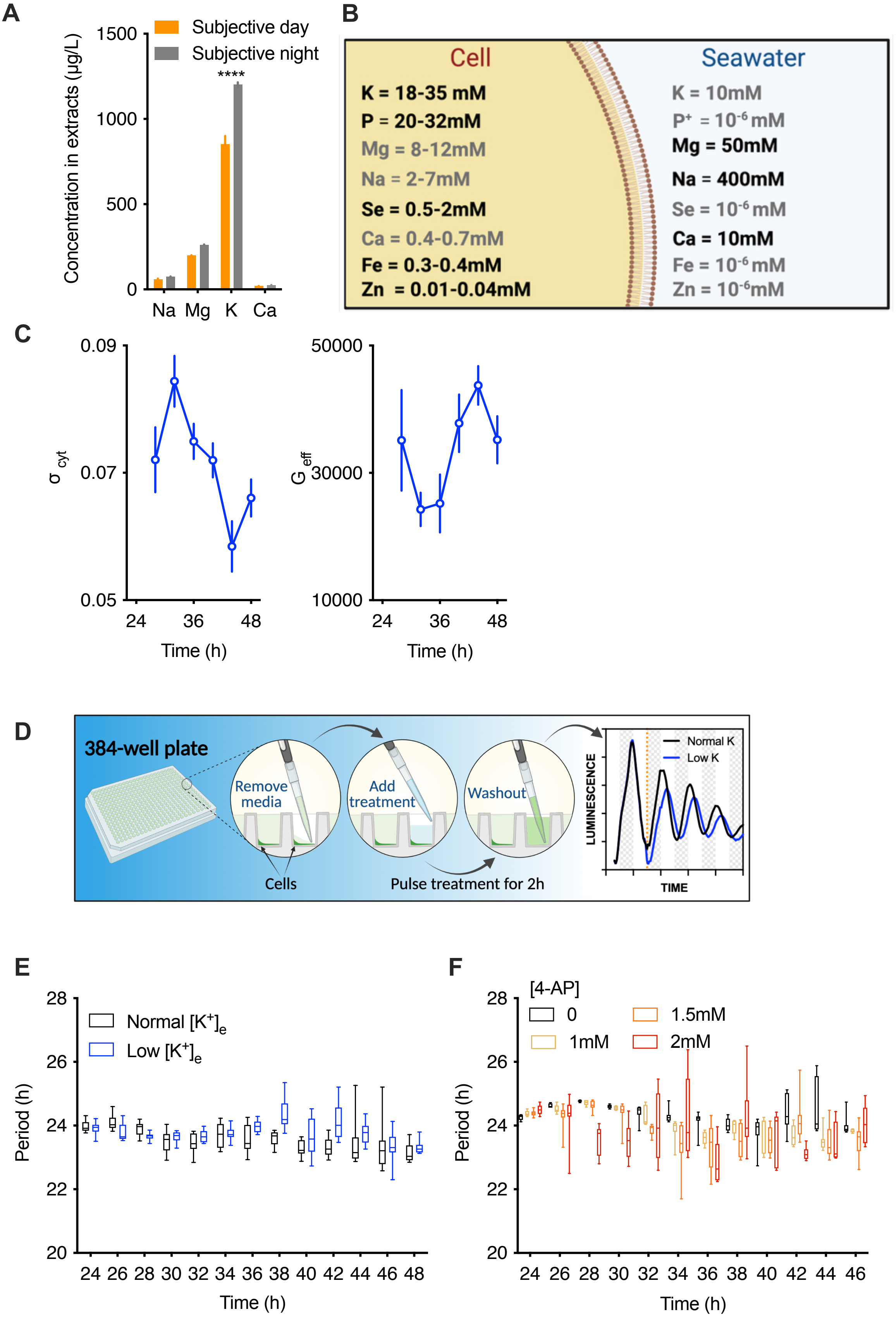


**Extended Data Figure 1**

### Extended Data Figure 2

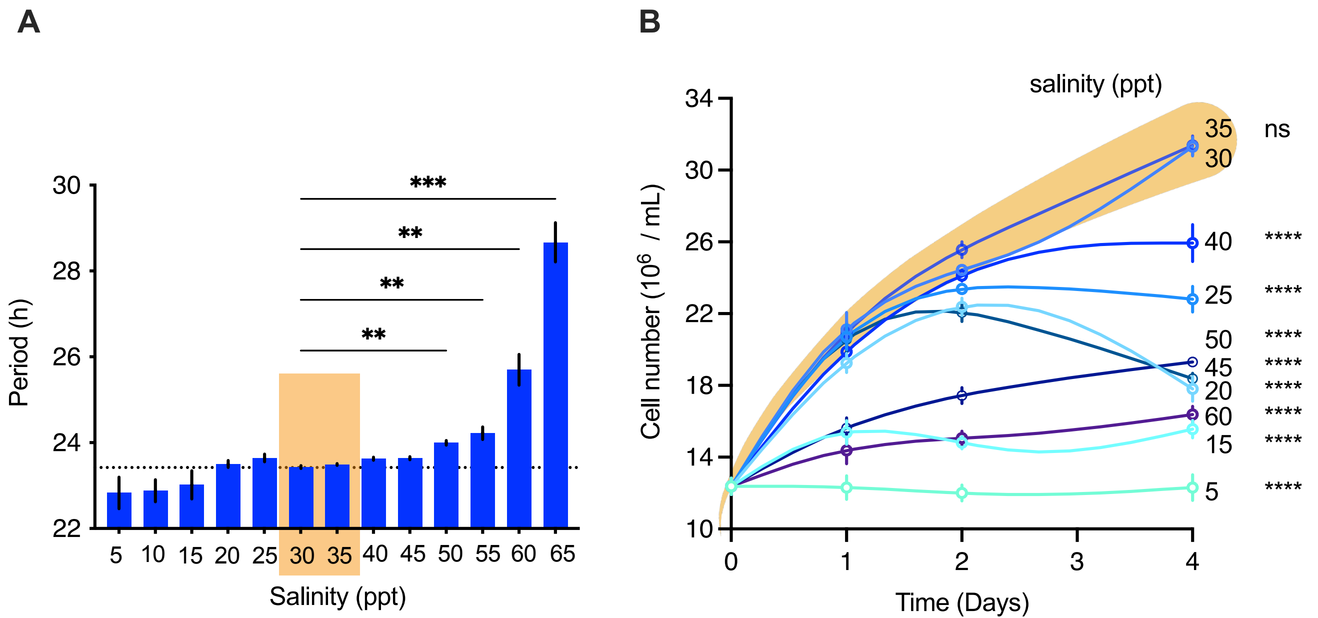


**Extended Data Figure 2**

### Extended Data Figure 3

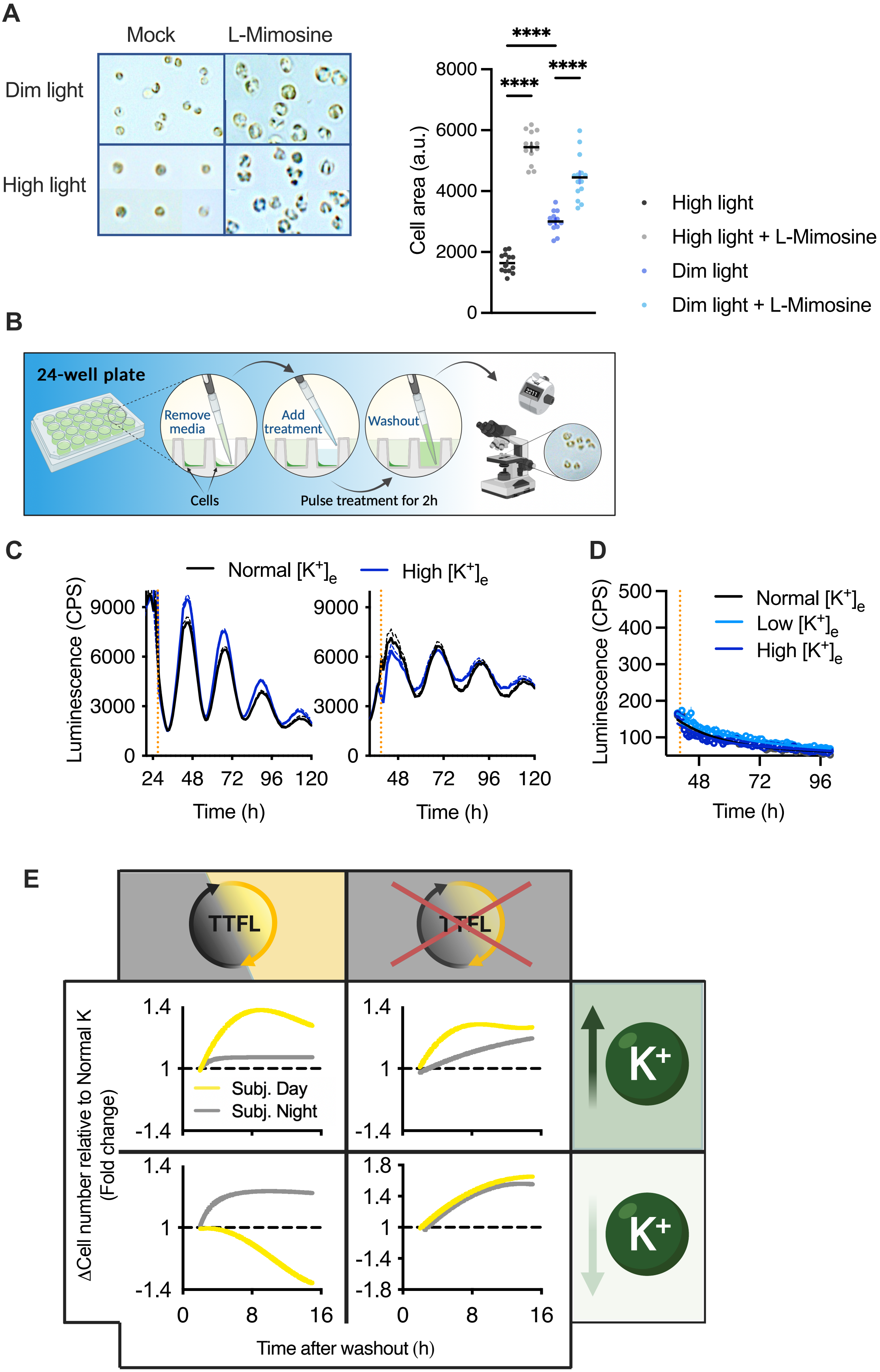


**Extended Data Figure 3**

### Extended Data Figure 4

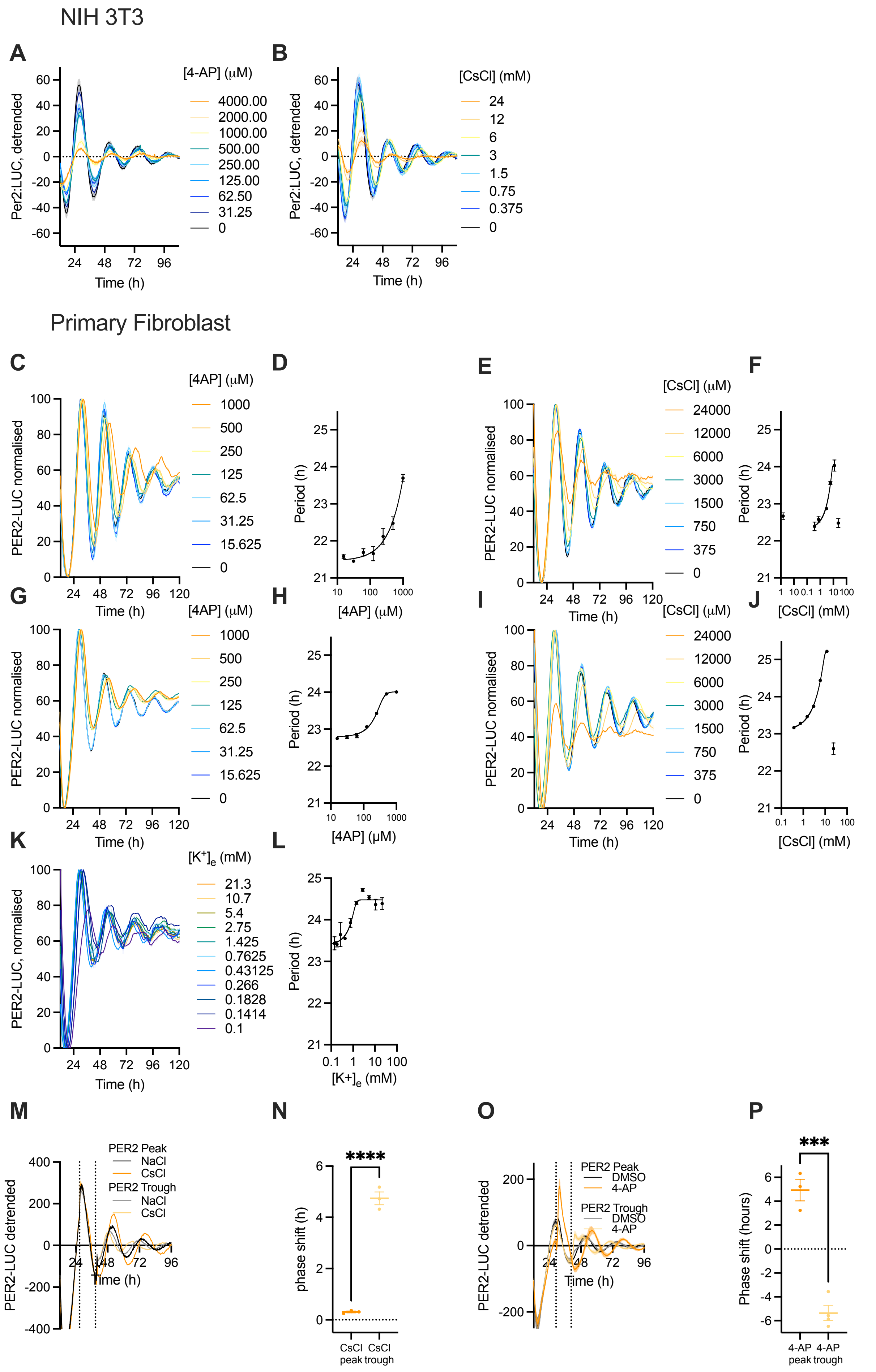


**Extended Data Figure 4**

### Extended Data Figure 5

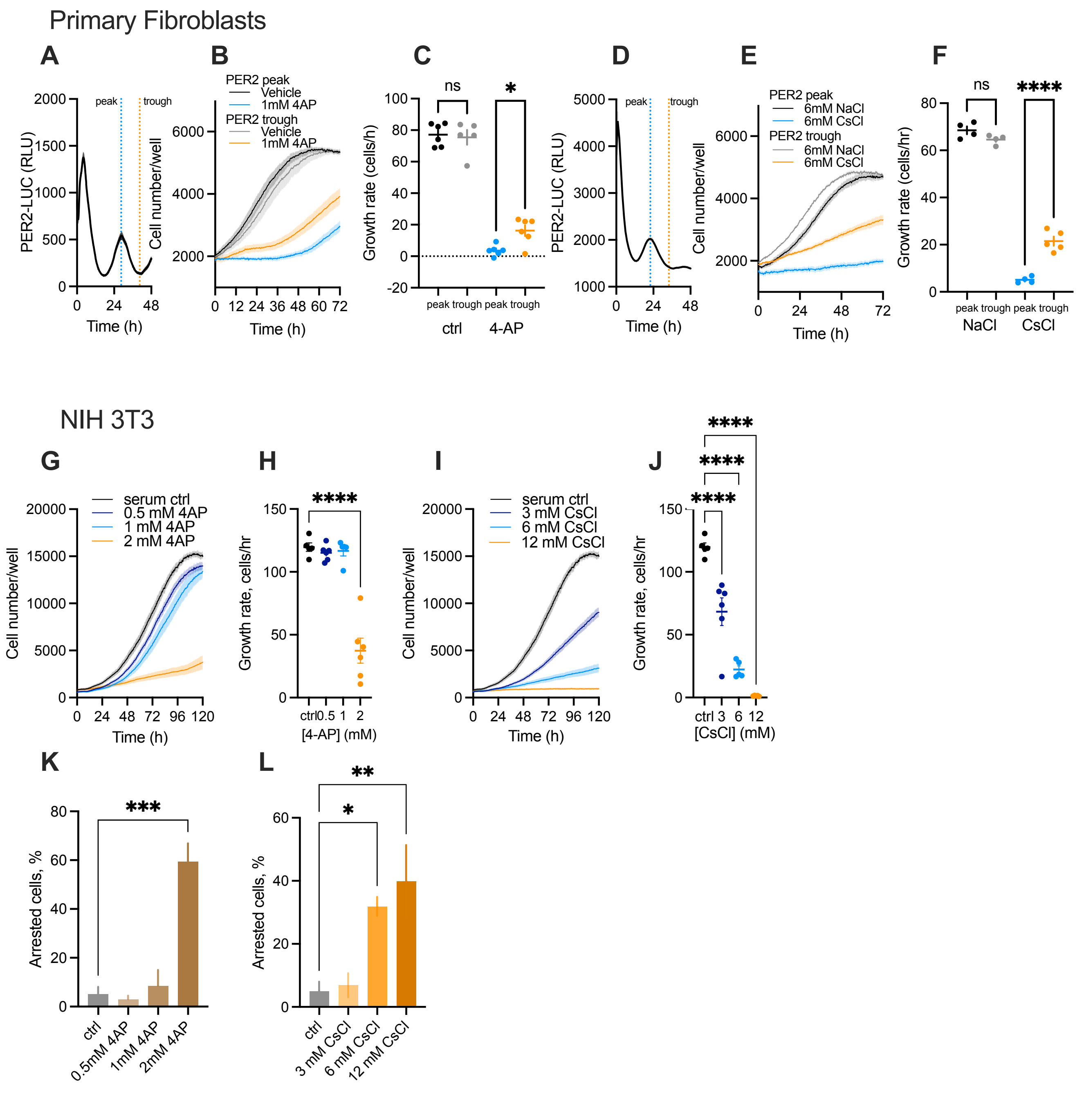


**Extended Data Figure 5**
